# Charting shared developmental trajectories of cortical thickness and structural connectivity in childhood and adolescence

**DOI:** 10.1101/572552

**Authors:** G. Ball, R. Beare, M. L. Seal

## Abstract

The cortex is organised into broadly hierarchical functional systems with distinct neuroanatomical characteristics reflected by macroscopic measures of cortical morphology. Diffusion-weighted MRI allows the delineation of areal connectivity, changes to which reflect the ongoing maturation of white matter tracts. These developmental processes are intrinsically linked with timing coincident with the development of cognitive function.

In this study, we use a data-driven multivariate approach, non-negative matrix factorisation, to define cortical regions that co-vary together across a large paediatric cohort (n=456) and are associated with specific subnetworks of cortical connectivity.

We find that age between 3 and 21 years is associated with accelerated cortical thinning in fronto-parietal regions, whereas relative thinning of primary motor and sensory regions is slower. Together, the subject-specific weights of the derived set of components can be combined to predict chronological age. Structural connectivity networks reveal a relative increase in strength in connection within, as opposed to between hemispheres that vary in line with cortical changes. We confirm our findings in an independent sample.

## Introduction

Brain development during childhood and adolescence has been well-characterised in recent years using Magnetic Resonance Imaging (MRI) (Mills et al. 2016). MRI provides a non-invasive method to examine neuroanatomy at a multiple scales. Analyses of structural MRI have found that the volume of grey and white matter follows different developmental trajectories during childhood.

Longitudinal analyses describe widespread and regionally variable decreases in cortical volume and thickness over childhood, with the largest decreases in thickness observed in the parietal lobe and the smallest changes in primary sensory-motor cortex (Wierenga et al. 2014; Vijayakumar et al. 2016; Tamnes et al. 2017). During childhood, observable patterns of cortical thinning, in part, reflect changes to the tissue microstructure (Huttenlocher 1979; Pakkenberg and Gundersen 1997; Vértes and Bullmore 2015). Over the same period, significant maturation of the brain’s white matter occurs, with increases in overall volume and continuing axonal myelination (Tamnes et al. 2010; Lebel and Beaulieu 2011; Lebel and Deoni 2018).

The cortex is organised into broadly hierarchical functional systems with distinct neuroanatomical characteristics, including neuronal density and cortico-cortical connectivity (Felleman and Van Essen 1991; Collins et al. 2010; Markov et al. 2013). Areal differences in cortical cytoarchitecture mirror this hierarchy (Collins et al. 2010), a pattern that is reflected by macroscopic measures of cortical morphology (Wagstyl et al. 2015). At a microscopic level, neuronal density is inversely related to cortical thickness (Schüz and Palm 1989; Cahalane et al. 2012; Wagstyl et al. 2015), whereas synaptic density tends to increase with increasing thickness (Schüz and Palm 1989).

Diffusion-weighted MRI allows the delineation of macroscale areal connectivity, changes to which reflect the ongoing maturation of white matter tracts. These developmental changes vary in timing and extent in a regional pattern that likely reflects the functional organisation of the brain. Studies using diffusion MRI have shown markers of tissue organisation, including fractional anisotropy, increase with age during childhood (Lebel et al. 2008), with maturation of commissural and projection fibres regions preceding intra-hemispheric association fibres (Lebel and Beaulieu 2011). The effects of white matter maturation on structural connectivity measures reveal large-scale topological organisation of the structural connectome with measures of network efficiency and modularity increasing over childhood and adolescence (Zhao et al. 2015; Baum et al. 2017).

It is likely these developmental processes are intrinsically linked (Van Essen 1997; Herculano-Houzel et al. 2010; Mota and Herculano-Houzel 2012) with timing coincident with the development of cognitive function (Blakemore and Choudhury 2006). In children, evidence of increased cortical thinning and white matter maturation from MRI is associated with improved cognitive performance (Kharitonova et al. 2013; Squeglia et al. 2013; Krogsrud et al. 2018).

Regional variation in both grey and white matter development reflects the functional organisation of the brain, with lower-order motor and sensory regions developing before areas that support complex, executive function. In a recent study, Sotiras et al. used a data-driven, multivariate method for dimension reduction, non-negative matrix factorisation (NMF), to identify a set of regional patterns of coordinated decreases in cortical thickness during adolescence (Sotiras et al. 2017). They found that changes in cortical thickness were spatially heterogeneous, with regional variation mirroring functional organisation and genetic patterning (Sotiras et al. 2017). This modular organisation is supported by evidence that cortical regions can be defined based on the extent of their shared anatomical connections (Hilgetag et al. 2000; Bullmore and Sporns 2012). Both neuronal density and dendritic spine density are correlated to areal connectivity (Scholtens et al. 2014; Beul and Hilgetag 2019) and white matter networks derived from MRI can similarly be split in subnetworks of connections, or modules, that appear to support known cortical systems, vary together with age and can be disrupted in neurodevelopmental disorder (Ball, Beare, et al. 2017; Baum et al. 2017; Faskowitz et al. 2018). Therefore, macroscopic markers of brain development, including cortical thickness and measures of white matter connectivity reflect regionally-varying patterns of complex, interlinked microscopic processes across the white and grey matter.

Multivariate methods, including NMF, are well-suited to neuroimaging analysis, enabling pattern discovery in large, complex datasets (Beckmann and Smith 2005; Calhoun et al. 2009; Sotiras et al. 2015; Ball, Beare, et al. 2017). Recently, a number of multivariate methods have been described that aim to integrate measures from different imaging modalities (Groves et al. 2011; Sui et al. 2012). This allows correlated patterns to be identified across imaging types, providing multiple views of developmental processes across tissue compartments. NMF is particularly apt for MRI analysis given that most MR-derived anatomical measures are non-negative (cortical thickness, tissue volume, fractional anisotropy, etc.) (Sotiras et al. 2015). When applied to anatomical data, NMF returns sparse components that represent local patterns, or components, of covariation, that permit interpretation as regions of coordinated growth or maturation (Sotiras et al. 2015, 2017). We have also previously applied this method to structural network data, revealing subnetworks, or network components, with connections that vary together across subjects (Ball, Beare, et al. 2017).

In this paper, we aim to combine these approaches to describe how patterns of regional change in cortical thickness are associated with specific patterns of change to structural connectivity networks during childhood and adolescence. In a large development cohort, we demonstrate that NMF provides hierarchical decomposition of the cortex based on regional differences in cortical thickness change over time. Additionally, using a supervised NMF approach we define specific patterns of structural connectivity that vary over the population in line with each cortical pattern.

## Methods

### Data

Data were acquired from the PING Study.(Jernigan et al. 2016) The PING cohort comprises a large, typically-developing paediatric population with participants from several US sites included across a wide age and socioeconomic range. The human research protections programs and institutional review boards at all institutions participating in the PING study approved all experimental and consenting procedures, and all methods were performed in accordance with the relevant guidelines, regulations and PING data use agreement (Brown et al. 2012). Written informed consent was obtained for all PING participants. Exclusion criteria included: a) neurological disorders; b) history of head trauma; c) preterm birth (less than 36 weeks); d) diagnosis of an autism spectrum disorder, bipolar disorder, schizophrenia, or mental retardation; e) pregnancy; and f) daily illicit drug use by the mother for more than one trimester.(Jernigan et al. 2016) Similar proportions of males and females participated across the entire age range. PING data are available via the NIMH Data Archive (ID 2607).

The PING cohort included 1493 participants aged 3 to 21 years, of whom 1249 also had neuroimaging data. Of these, n=763 imaging datasets were available to download.

### Neuroimaging

At each site, pulse sequences were optimized for equivalence in contrast properties across scanner manufacturers (GE, Siemens, and Phillips) and models (Brown et al. 2012; Jernigan et al. 2016). T1-weighted images were acquired using standardized 3-Tesla high-resolution 3D RF-spoiled gradient echo sequences with prospective motion correction (PROMO). Diffusion data were acquired using axial 2D EPI scans with 30-directions, *b*-value = 1000 and slice thickness = 2.5mm with a corresponding reverse-encoded *b* = 0 map for EPI *B*_0_-distortion correction. Detailed acquisition parameters for each scanner manufacturer can be viewed at: http://pingstudy.ucsd.edu/resources/neuroimaging-cores.html

Quality control for the PING data is detailed in Jernigan et al. (2016). In brief, images were inspected for excessive distortion, operator compliance, or scanner malfunction. We performed additional, on-site, visual quality control assessment of T1 and diffusion data. This involved a visual inspection of T1 volumes for motion or image acquisition or reconstruction artefacts. In addition, we included only diffusion data with at least one complete 30-direction acquisition, after inspection for slice-dropouts or motion artefacts. Five participants were removed due to excessive motion in T1-weighted images; a further 312 were removed due to a lack of matched 30-direction sequences (n=232), or due to motion or image artefacts (n=80).

This resulted in a final cohort of n=456 participants with both T1-weighted and 30-direction diffusion data from 5 participating sites (mean age [range] = 12.6 [3.2-21.0] y, 233 male [51.1%]). Site-specific demographic data are shown in Table S1.

### Discovery and validation cohorts

The full cohort was split into discovery and validation cohorts prior to analysis. A random subset of 152 subjects (33%) were removed and retained as a held-out validation cohort.

### Image processing

T1 MRI data was processed as in Ball et al (Ball, Adamson, et al. 2017). Briefly, vertex-wise maps of cortical thickness were constructed with FreeSurfer 5.3 (http://surfer.nmr.mgh.harvard.edu). This process includes removal of non-brain tissue, transformation to Talairach space, intensity normalisation, tissue segmentation and tessellation of the grey matter/white matter boundary followed by automated topology correction. Cortical geometry was matched across individual surfaces using spherical registration.(Dale et al. 1999; Fischl et al. 1999, 2002; Fischl and Dale 2000) Any images that failed initial surface reconstruction, or returned surfaces with topological errors, were manually fixed using white matter mask editing and re-submitted to FreeSurfer until all datasets passed inspection. To reduce computational load, data were downsampled to the ***fsaverage5*** surface comprising 10242 vertices per hemisphere.

Diffusion data were pre-processed using MRtrix 3.0 (http://www.mrtrix.org/). Data were first denoised and corrected for EPI distortions using ***topup/eddy*** (Andersson and Sotiropoulos 2016; Veraart et al. 2016). The diffusion signal at each voxel was then modelled using a tensor model, from which we calculated fractional anisotropy (FA, Basser and Pierpaoli 1996).

### Whole-brain tractography

Prior to tractography, each T1 image was segmented into 3 tissue classes (grey matter, white matter, cerebrospinal fluid) using FSL’s FAST (Zhang et al. 2001) and the cortical grey matter parcellated into n=220 cortical regions-of-interest derived from subdivisisions of the Desikan-Killiany anatomical atlas using ***easy_lausanne*** (https://github.com/mattcieslak/easy_lausanne) (Hagmann et al. 2008; Daducci et al. 2012). Tissue segmentations and parcellations were transformed into diffusion space using Boundary-Based Registration (Greve and Fischl 2009) followed by nonlinear registration using ***ANTS*** (Avants et al. 2008).

Probabilistic whole-brain fibre-tracking was performed using wild bootstrap diffusion tensor tractography (Jones 2008) implemented in MRtrix3. Ten million streamlines were generated per participant with a step size of 1mm. Streamlines were seeded from the grey/white matter interface and anatomically constrained using a 5-tissue-type (5TT) mask (Smith et al. 2012).

Structural connectivity matrices (of size 220 × 220) were constructed by identifying streamlines that connected cortical regions-of-interest using a 2mm radial search from streamline endpoints. For each pair of connected regions, connection strength was estimated by calculating mean FA along connecting streamlines.

Connectivity matrices were subject to a consistency-based thresholding, retaining the top 10% most consistent edges across subjects (n=1538) (Roberts et al. 2017).

### Site correction

To correct for possible effects of acquisition site on the initial NMF decomposition, we used ComBat harmonisation (https://github.com/ncullen93/neuroCombat) to estimate and remove linear site effects from vertexwise measures of cortical thickness (see also Fig S3). This method has been shown to remove unwanted sources of scan variability in in multisite analyses of cortical morphometry measures (Fortin et al. 2018).

### Projective non-negative matrix factorisation

We decomposed cortical thickness data into a set of non-negative, orthogonal components using projective NMF (Yang and Oja 2010; Sotiras et al. 2015; Ball, Beare, et al. 2017; Sotiras et al. 2017).

Briefly, non-negative matrix factorisation (NMF) is a multivariate method that models an n feature × *m* sample data matrix, V, as the product of two non-negative matrices: W with dimensions *n* × *r* and H with dimensions *r* × *m*, such that *V* ≈ *WH*. Generally, the number of basis components *r* < min(*m, n*), thus *WH* represents a low-rank approximation of the original data in *V* (Lee and Seung 1999).

In projective NMF, the matrix, H is replaced with W^T^V such that *V* ≈ *WW*^*T*^*V*. PNMF confers a number of benefits over standard NMF including fewer learned parameters, and increased sparsity of the resulting matrix *W* (Yang and Oja 2010).

In this context, the columns of *W* represent a set of components, each comprising cortical regions that co-vary together across the study population and *W*^*T*^*V* provides a corresponding set of subject-specific weights, one per component (Sotiras et al. 2015). Together, these elements can be combined to approximately reconstruct any subject’s original dataset.

We implemented PNMF in Python (3.6.3), performing 20,000 iterations each time. Before NMF, each subject’s cortical data were normalised to unit norm as an equivalent operation to correcting each vertex for mean cortical thickness, while ensuring non-negativity.

### Supervised NMF

To identify structural connections that co-vary with each of the cortical components, we performed a supervised NMF decomposition of the structural connectivity data. We derive the subject loadings from the PNMF decomposition of the cortical data: 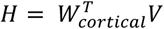 and decompose the connectivity data, V_conn_ by iteratively updating W_conn_ to minimise the (Euclidean) distance between the original and reconstructed matrices, subject to non-negativity constraints, while ensuring *H* remains fixed:

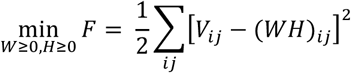

This results in a set of orthogonal network components, each comprising a sparse set of (topologically) localised connections (i.e.: the edge structure of different components does not overlap) and each varying across the population in line with a given cortical component. As with the cortical data, each subject’s connectivity was normalised prior to NMF.

### Reconstruction error, cross-validation and age prediction

For each set of components, we calculated the root mean square error (RMSE) between the original data matrices and matrices reconstructed from the NMF components for both cortical thickness and structural connectivity. To reduce bias in our error estimation, we employed Wold holdouts (Wold 1978), masking a random subset of 20% elements from each data matrix during NMF estimation and calculating error within the held-out subset. Holdouts were repeated five times and reconstruction error averaged across repeats.

In addition, to determine how well the low-rank data representations capture biological variability, we used each set of component timecourses to predict chronological age. We used Gaussian process regression implemented in ***scikit-learn*** (v0.20.0) to estimate age for each subject based on their individual component weights within a 10-fold cross-validation framework (Cole et al. 2017). For each set of components, age prediction error was calculated as the mean absolute difference between predicted and chronological age.

### Modelling component timecourses

For each component, we used Generalised Additive Models (GAMs) to model the relationship between age and component weight using the ***mgcv*** package in ***R***v3.5.1 (Wood 2017). GAMs are flexible models that allow a data-driven estimation of (possibly nonlinear) relationships between variables. Component weight was modelled as a smooth function of age, estimated using penalised thin plate regression splines with automatic smoothness estimation maximising marginal likelihood (ML) (Wood 2003, 2011). For each component, we additionally tested the inclusion of sex and site effects alongside age and compared model fit using the Akaike Information Criterion (AIC).

### Hierarchical decomposition

Hierarchical clustering was performed using Ward’s linkage based on Euclidean distance between the component timecourses. The cophenetic correlation was calculated between pairwise similarity and dendrogram distance of component timecourses as a measure of hierarchical organisation. Statistical significance was determined using random permutations (p<0.05, n=10000).

## Results

### Reconstruction error and age prediction

In the discovery cohort, we performing projective NMF to decompose cortical thickness data into 2, 5 10, 15 and 20 components and used the resulting component timecourses to derive a corresponding set of structural connectivity components.

To determine how well the full thickness and connectivity datasets were represented by each set of NMF components, we calculated reconstruction error at each level using five repeated Wold holdouts (Figure 1). As expected, training error decreased with increasing number of NMF components for both cortical thickness and structural connectivity. In contrast, reconstruction error of held-out data points increased moderately for cortical thickness (mean RMSE for 2, 5, 10, 15, 20 components: 0.001634, 0.001636, 0.001638, 0.001640, 0.001641, respectively). For structural connectively, test error remained relatively stable across component sets, decreasing slightly from 0.00766 for 2 components to 0.007652 for 20 components.

**Figure 1:**
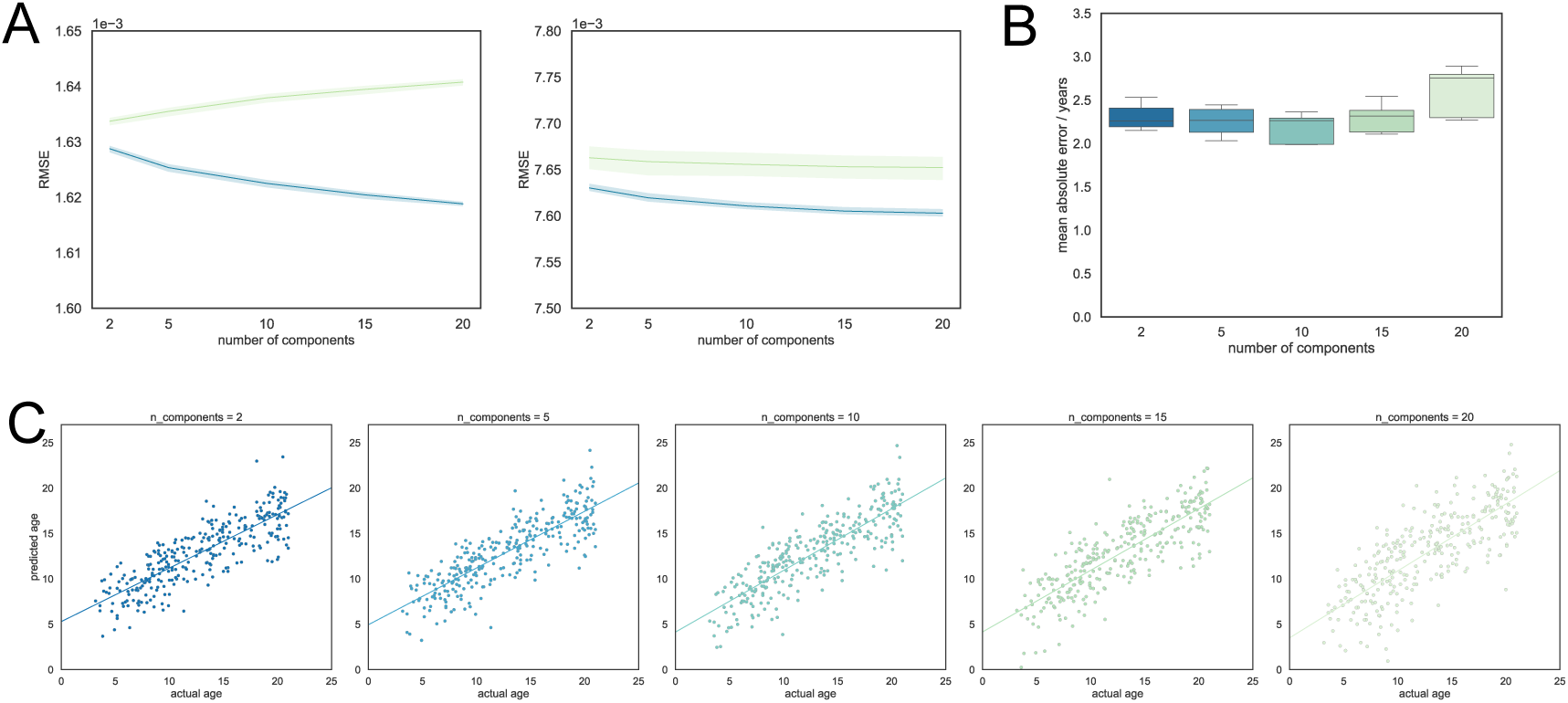
Reconstruction error and age prediction for increasing number of NMF components. A) reconstruction errors, average over 5 Wold holdouts, for cortical thickness (left) and structural connectivity data (right). Error for training datapoints (blue) and held-out test datapoints (green) are shown with 95% C.I. B) mean absolute error in age prediction is shown, averaged over 10 cross-validation folds for each set of NMF components. C) Individual age predictions are shown for each set of components.

To examine how well biological variability was captured by each set of components, we used Gaussian process regression to predict age from the set of component timecourses at each level. Prediction error was estimated using 10-fold cross-validation within the discovery cohort (Figure 1B). Figure 1C shows individual, cross-validated predictions of age based on each of the NMF component sets. Mean absolute error in age estimation was similar across all component sets with the lowest error observed in the 10 component set (mean MAE ± S.D = 2.18 ± 0.18) and highest in the 20 component set (2.60 ± 0.30).

As reconstruction errors and age prediction were relatively similar between component sets, we initially progress by focusing on the five component set only. For comparison, decompositions of cortical thickness and structural connectivity into 2, 5 and 10 components are shown in Fig S1.

### Cortical thickness components

Figure 2 shows the result of projective NMF decomposition of cortical thickness data into five components, along with corresponding component timecourses. Each image represents the spatial distribution of component weights across the cortex, highlighting regions that vary across the population together. Each map is associated with a timecourse that describes the relative contribution of that component to the full dataset and reflects how cortical thickness within the regions described by each map varies over time.

**Figure 2:**
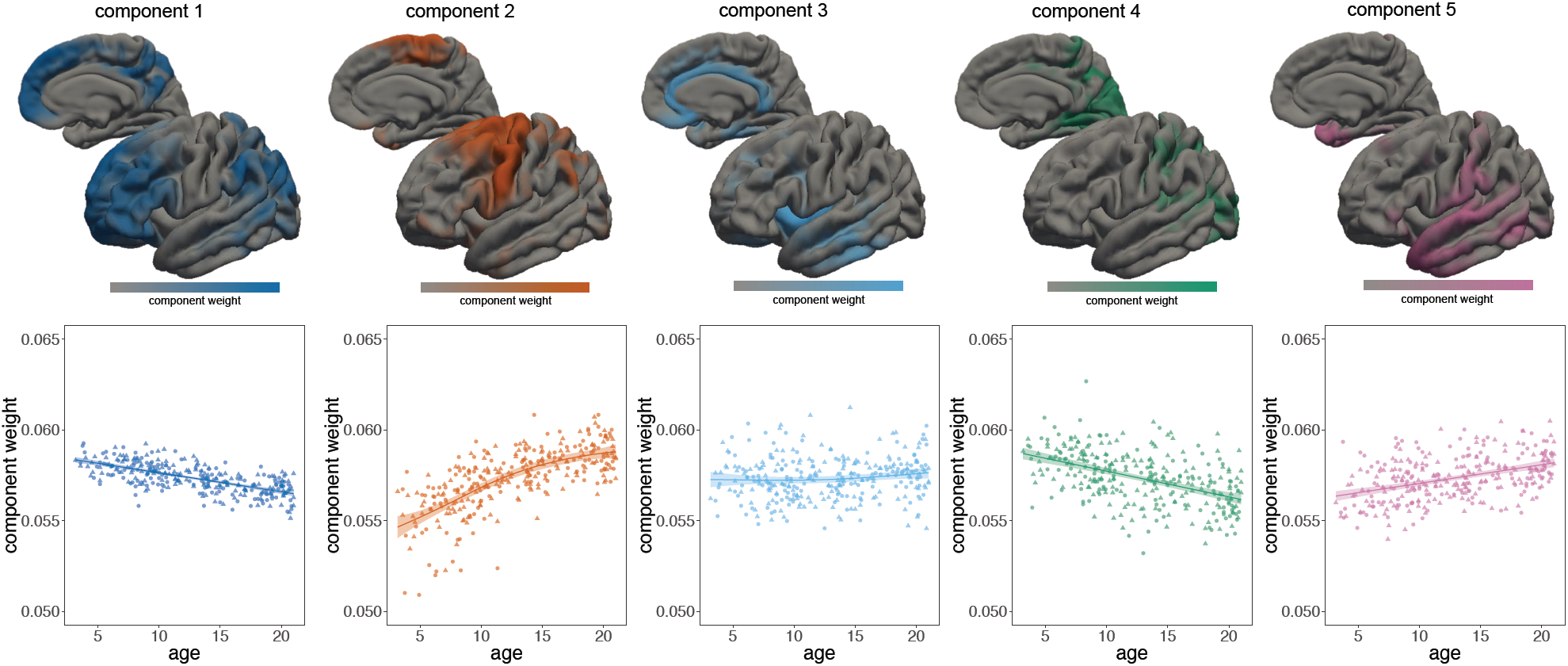
Cortical NMF components. Spatial maps for the five-component solution are shown with associated timecourses. Components weights are normalised to account for the mean trend in cortical thickness over time.

Component maps appear to group together functional subsystems including fronto-parietal networks (component 1), primary and supplemental motor regions (component 2), sensory regions (components 4 and 5) and cortical limbic systems (components 3).

As each subject’s data were normalised to a unit norm prior to NMF decomposition, the component timecourses in Figure 2 are effectively corrected for mean trends over time. We find that overall, mean cortical thickness decreased between 3 and 21 years (Figure S2). Thus, the timecourses presented in Figure 2 show changes relative to this overall trend and a decrease over time represents a regional decrease that is faster than the mean trend and ***vice versa***. The effect of this normalisation is shown in Figure S2.

Component 1 (dark blue, Figure 2) groups together dorsolateral and medial frontal cortex with lateral parietal cortex and the precuneus. Together, cortical thickness in these regions decreases between 3 and 21 years of age. In contrast, component 2 groups together primary and supplementary motor cortex, which ***relative to the mean trend***, increases over the same age span. Component 3 primarily includes insular and cingulate cortex and remains relatively stable over time. The final two components comprise primary visual cortex (component 4) and primary sensory along with superior temporal cortex and the anterior temporal horn (component 5), regions that increase (comp. 4) and decrease (comp. 5) relative to the mean trend over time. There were no significant differences between male and female timecourses for any component. Similarly, after applying ComBat harmonisation to the cortical thickness data prior to NMF, there were no significant site effects on component weight (Figure S3).

### Structural connectivity components

The timecourses derived from the initial projective NMF were used to drive a supervised decomposition of structural connectivity data to identify groups of connections that share the same relative trajectory over the population as each of the cortical components. As with the cortical data, connectivity data were normalised prior to decomposition. The mean trend in edge strength, indexed by fractional anisotropy, followed an upward course, increasing rapidly to around 12 years before plateauing. The mean trend and raw (unnormalised) component timecourses are shown in Figure S2. Figure 3 shows the corresponding structural connectivity subnetworks for each cortical component in Figure 2. The top 25% of edges based on component weight are shown for each subnetwork.

**Figure 3:**
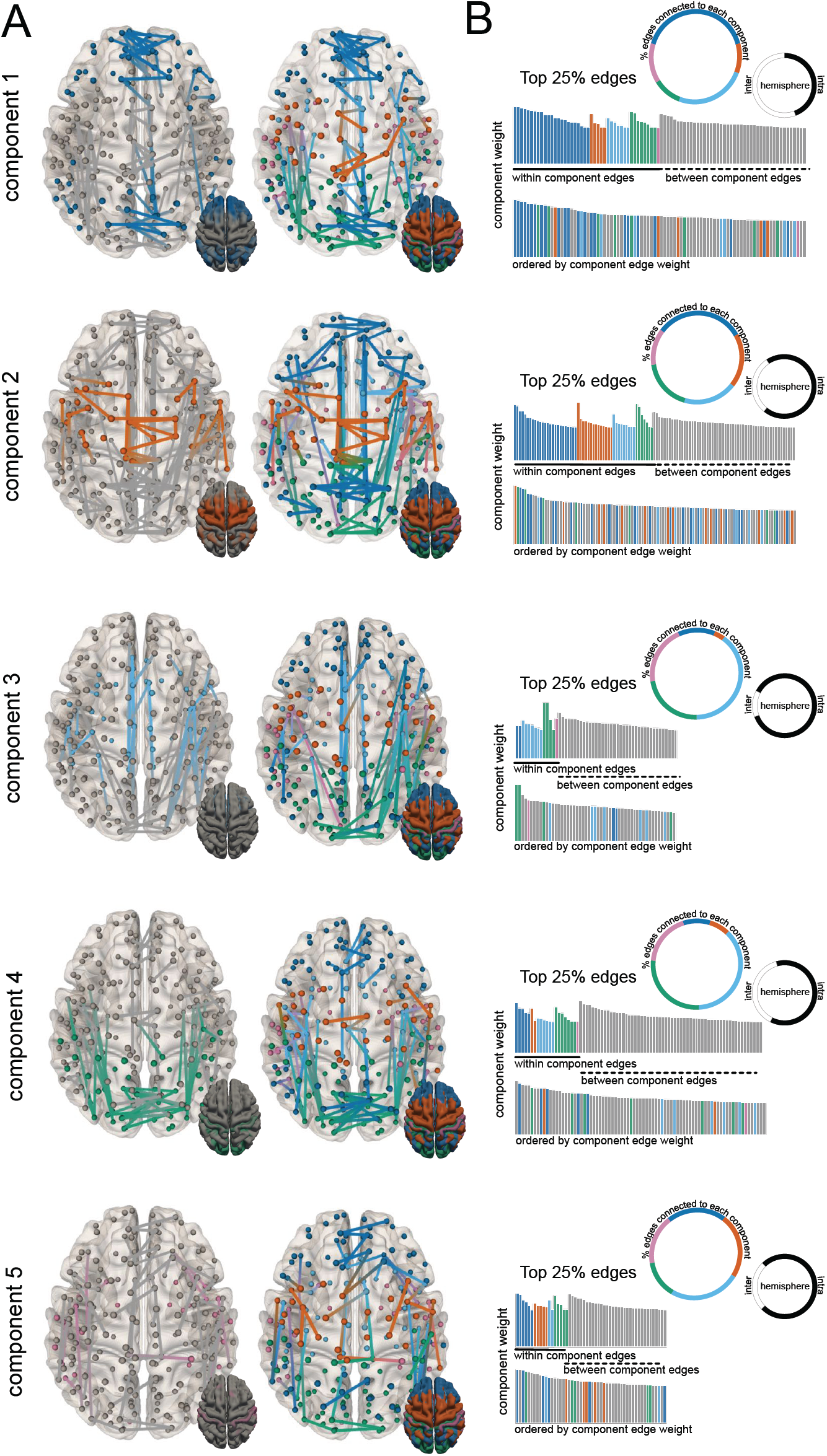
Connectivity NMF components. A) Connectivity subnetworks are shown in the same order as corresponding cortical components. The top 25% edges based on component weight are coloured based on connections to the corresponding component (left) or all components (right). B) Networks are summarised based on the proportion of edges connected different modules (coloured circle), the proportion of inter- and intra-hemispheric edges (black/white circle) and as a histogram of edge weights ordered by within- or between-modules connectivity and edge strength.

The NMF decomposition resulted in five distinct connectivity subnetworks, each of which contained connections both within and between the corresponding cortical components. Connectivity components 1 and 2 comprised approximately 50% within- and 50% between-component edges (Figure 3B). In contrast, between-component edges formed the majority of connections in connectivity components 3 (74%), 4 (75%) and 5 (64%).

Connectivity component 1 contained a large proportion (∼55%) of interhemispheric connections between homologous regions in medial frontal cortex (Figure 3A, 3B). While connections are present between all cortical components, edges within the corresponding cortical component formed the top ∼10% when ordered by component weight (dark blue, Figure 3B). Additional connections are seen between homologous regions in primary visual and motor cortex (cortical components 2 and 4) and intra-hemisphere connections between insular and ventral parietal cortex (components 3 and 4). As shown by the timecourses in Figure 2, the relative contribution of this subnetwork to the full connectivity dataset changes at a slower rate that the mean trend, with component weight (along with component 4) higher relative to other components at 3 years of age, and lowest at 21 years (Figure S2).

In contrast, edges forming the second connectivity component follow a similar distribution of intra- and inter-component connections (Fig 3A), but the majority of edges connected intra-hemispheric regions and component weight increased rapidly over the age range resulting in the largest relative contribution to the full network at the end of the age range (Fig S2). Unlike connectivity component 1, the majority of the strongest edges do not connect primarily within regions of the corresponding cortical component. Connections are present between frontal and parietal regions (cortical component 1) and between hemispheres in supplemental motor (component 3), frontal, parietal (component 1) and visual cortex (component 4)

Connectivity component 3 is represented largely by intra-hemispheric connections (87%) between different cortical components, with a large proportion of edges connecting one or more nodes in cortical component 3 or 4 (Fig 3B). As with cortical component 3, this relative strength of this subnetwork increases over time with the mean trend (Fig S2). This reciprocal connectivity is also evident in connectivity component 4, with 64% of edges connecting to a node in either cortical component 3 or 4. The fourth subnetwork also displays a pattern of extensive inter-hemispheric connections between occipital regions within the corresponding cortical component (Fig 3A).

Connectivity component 5 has contains no edges that connect solely within the corresponding cortical component. Instead connections are evenly distributed between cortical components, with the majority of edges connecting regions within hemisphere (Fig 3B). As with connectivity component 2, component weight increases relative to the mean trend over the age range.

### Component hierarchy

To determine whether NMF provides a hierarchical decomposition, i.e.: by splitting larger components into smaller constituent parts. We performed projective NMF on the cortical thickness data, specifying 20 components and clustered the cortical maps and corresponding structural connectivity networks according to a hierarchical clustering of the component timecourses (Figure 4). There was a significant correlation between timecourse similarity over the 20 components and distance along the clustering dendrogram (cophenetic correlation= 0.66, p<0.001 10,000 random permutations). By combining maps and networks from the higher level decomposition according to the hierarchical clustering algorithm, we were able to recapture patterns derived from NMF decompositions at coarser levels (2, 5 and 10 components (Figure 4B and C) demonstrating a stable, hierarchical decomposition of both datasets.

**Figure 4:**
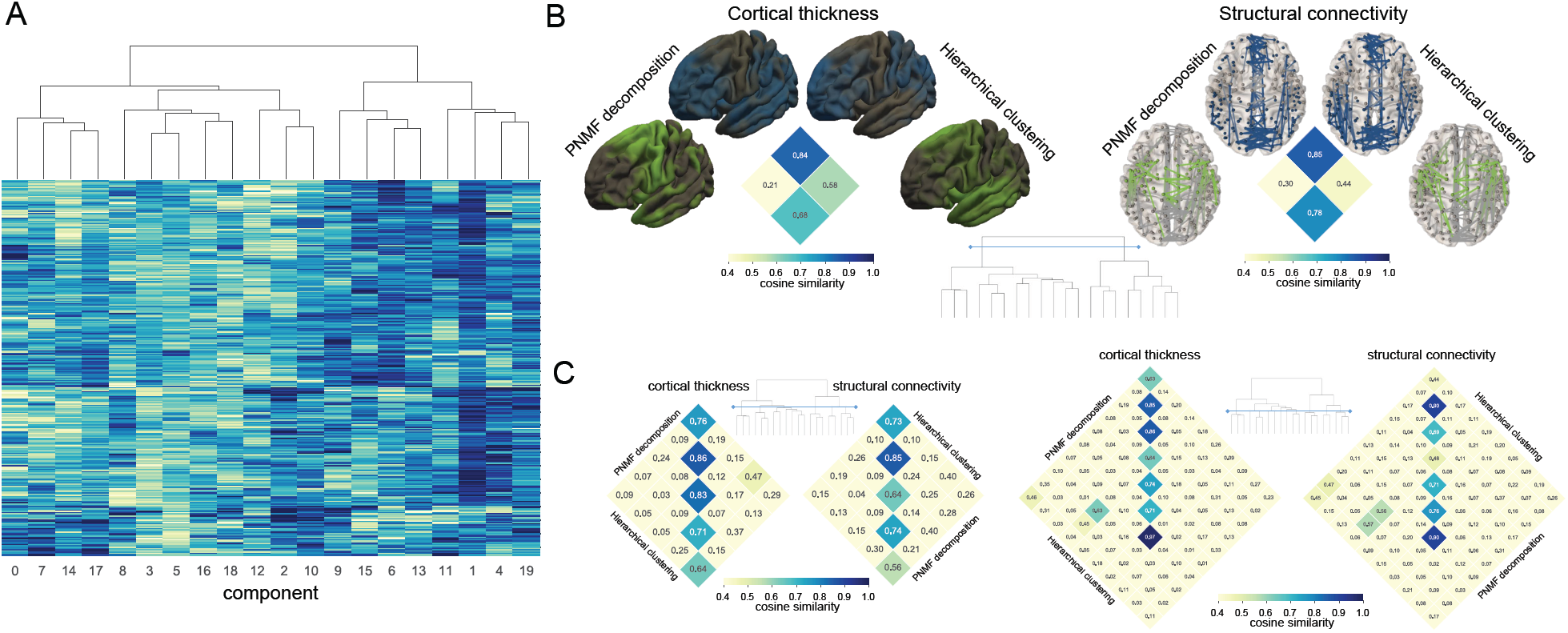
Hierarchical decomposition of NMF components. A) 20 NMF components are clustered based on similarity of component weights across subjects. B) The spatial similarity between NMF decompositions at a level of 2 components are shown with maps constructed through the addition of hierarchically clustered components at lower levels. C) Similarity matrices are shown at the 5 and 10 component level.

### Validation cohort

For validation, we repeated the NMF decomposition of cortical thickness and structural connectivity data into five components in the remaining, held-out subsample (n=152). The similarity of the resultant cortical maps and connectivity subnetworks are shown in Figure 5, ordered by average similarity to matched components in the discovery cohort. Average cosine similarity ranged from 0.4 (component 5) to 0.57 (component 2). Matching was performed separately for cortical and connectivity components; similarity was higher for cortical thickness maps (range: 0.53 – 0.64) compared to connectivity subnetworks (0.27 – 0.50).

**Figure 5:**
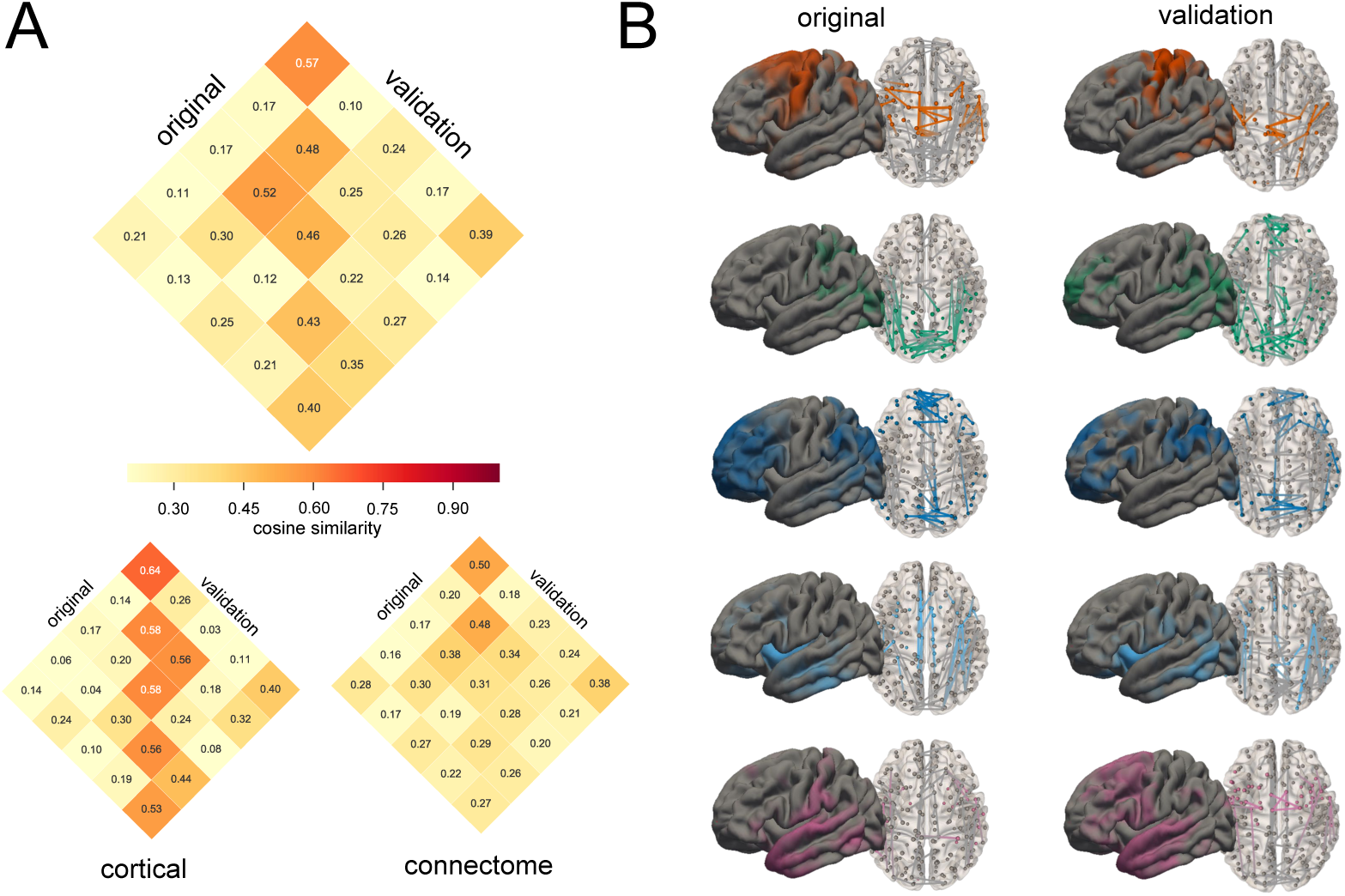
Comparison of discovery and validation cohorts. A) The average similarity between matched components in the original and validation cohorts are shown (top), within corresponding matrices for similarity of cortical maps and connectivity networks. B) Comparison of cortical and connectivity components from the original and validation cohorts, ordered by average similarity.

## Discussion

In this study, we employ data-driven, matrix factorisation methods to decompose the developing cortex into a set of modules containing areas that co-vary together with age and are correlated with specific structural connectivity subnetworks.

We find that increasing age between 3 and 21 years is associated with accelerated thinning of fronto-parietal regions, whereas relative thinning of primary motor and sensory regions is decreased. Together, the weights of each set of components can be combined to predict chronological age and can be decomposed further into sets of smaller sub-modules while retaining overall data structure. Through the use of a second, supervised decomposition, we define a set of areal connections, with an overall weight that varies across the population according to the same trajectory as each of the cortical components. From this, we determine that connections between cortical areas within the same hemisphere increase in strength with age faster than connections between hemispheres, as well as observing patterns of differential maturation in connections within and between cortical modules.

Previously, Sotiras et al used NMF to demonstrate that the cortex can be split into a set of components, or modules, that co-vary together, capture developmental change over time and reflect the organisation of large scale functional networks (Sotiras et al. 2017). Here, we replicate these findings in a separate cohort, extending their observations to test the hierarchical properties of NMF decomposition as well as combining this approach with diffusion MRI to map out co-varying patterns of structural connectivity. Overall, our findings support that of Sotiras et al.; we define a set of cortical components with spatial patterns that reflect known functional systems including: fronto-parietal, primary motor, sensory and cortical limbic systems (Yeo et al. 2011; Sotiras et al. 2017). These patterns replicate well in an independent subset of the cohort. We find an overall decrease in cortical thickness with age (Fig S2), this support recent findings in multiple longitudinal cohorts demonstrating a broadly monotonic decrease in thickness over time in the same age range (Tamnes et al. 2017). After correcting for this mean trend, we observe regional differences in rate of change in thickness across cortical components. In contrast to Sotiras et al., where raw (not normalised) cortical data were used for NMF, no differences were observed between sexes in component trajectory, suggesting any sex differences were captured by the differences in mean thickness. Focusing on the five component model, we find that cortical thinning was fastest in components 1 and 4 comprising regions in dorsolateral frontal and parietal cortex, precuneus and primary visual cortex, and slowest in primary and supplementary motor, primary sensory and superior temporal cortex. Regional modelling of cortical thickness has found similar patterns, broadly respecting the organisational hierarchy of the cortex (Tamnes et al. 2010, 2017; Fjell et al. 2015; Vijayakumar et al. 2016). We anticipate this difference in regional rate of change reflects the differential progress of microstructural processes including synaptic pruning (Huttenlocher 1979), although recent has evidence has suggests that apparent cortical thinning in the visual cortex may be dependent on tissue contrast changes due to intra-cortial myelination, rather than synaptic remodelling (Natu et al. 2018).

Our use of supervised NMF to derive a corresponding set of connectivity subnetworks that co-vary over the population in line with each cortical component allow the direct comparison of cortical and white matter development. Other studies have employed different methods to study concomitant changes across both tissue compartments. These approaches include inspection and comparison of white matter properties directly subjacent to different cortical regions (Croteau-Chonka et al. 2016), or examination of white matter tracts from specific cortical gyri (Jeon et al. 2015) or connections between cortical areas that displaying significant age-related changes in surface area in childhood (Cafiero et al. 2019). A strength of our approach is that we used a data-driven method to select important connections based on their co-variation with each cortical component. This allows the derivation of spatially (topologically) independent subnetworks, i.e.: where connections are not shared between components, that vary together across the population (Ball, Beare, et al. 2017). Importantly, this form of guided decomposition can be applied to any dataset, imaging or otherwise, shared by the same participants.

In contrast to previous approaches, this method does not restrict white matter connections to those that exclusively connect to the corresponding cortical component, instead selecting groups of edges where change over time relative to the mean trend mirrors that seen in the cortex. The mean trend in connectivity strength revealed a relatively quick increase up that slows at around 12 years reaching a plateau by late adolescence. Similar trends in white matter maturation, as indexed by FA have been described previously using tractographic approaches (Chen et al. 2016). Overall, connectivity component 2 showed the most rapid increases, with components 1 and 4 the slowest. Components 3 to 5 contain more between-component connections than within-, highlighting generalised patterns of maturation across the whole network. Connectivity component 1 exhibited the largest proportion of within-module connections between its respective cortical component regions, namely between homologous regions in frontal and parietal cortex. Edges within this component are predominantly inter-hemispheric and, as with the cortical component, the relative contribution of these edges to the full network decreases with age, rising slowest between 3 and 21 years. In contrast, connectivity component 2 contains edges that show the largest increase with age and connect predominately within hemisphere but are not limited to the corresponding cortical component, with edges shared between regions in each module. These findings support observations from diffusion tractography studies that found that inter-hemispheric commissural tracts display earlier maturation compared to within-hemisphere association fibres such as the superior longitudinal and fronto-occipital fasciculi (Lebel and Beaulieu 2011). A large proportion of reciprocal connectivity was observed between subnetworks associated with the spatially adjacent cortical components 3 and 4, with a number of edges connecting one areas in each component, although the components showed differential change over time. Component 4, comprising a larger portion of inter-hemispheric edges decreasing in strength relative to the mean compared to component 3. Component 5 revealed a relatively distributed subnetwork with edges connecting to all cortical components, predominately within hemisphere and with a strength that increased relatively quickly with age. Therefore, this component may represent an integrative network process that facilitates communication across the network and have been shown to increase during adolescence (Dennis et al. 2013).

As is often the case with unsupervised methods, the appropriate choice of component number poses a difficult challenge. We performed NMF using 5 different levels: 2, 5, 10, 15 and 20 and compared how well the resulting components could reconstruct the original data, and how well the resulting timecourses could be used to predict chronological age. Overall, we did not find significant differences between the different levels of resolution. Reconstruction error was relatively stable across all component sets, as was age prediction error for all levels except for n=20. Overall, age prediction error was in range with previous ‘brain age’ estimates (Franke et al. 2012; Ball, Adamson, et al. 2017; Cole et al. 2017) confirming that utility of NMF for providing useful low-rank representations of large imaging datasets (Varikuti et al. 2018).

As the prediction errors were relatively stable, we chose to focus on the five component set for ease of interpretation. We provide images of the cortical and network components in Fig S1 for comparison. We also note that, using split-half repeats as a measure of reliability, Sotiras et al. found the most reliable decompositions at similarly coarse levels (2, 7 and 18). The fact that reconstruction error did not significantly change when increasing the number of components suggested that the bulk of the original data matrix was captured by the lowest level of decomposition (i.e: 2 components), and that increasing the decomposition resolution further resulted in a subdivision of these two, larger components. To test this hypothesis, we performed a hierarchical clustering of the highest resolution decomposition, combining cortical and connectivity components based on the similarity of their timecourses. We found that there was significant hierarchical structure in the NMF decompositions, and could largely recapture the original 2-component solution by combining a number of smaller components (Figure 4). This may allow future studies to compare findings even if different numbers of components are specified.

Finally, we performed a within-cohort validation, running the NMF decompositions separately in a held-out subset of the full cohort. We found that, in general, similar spatial maps were defined in the validation set. Similarity was higher in the cortical components compared to the connectivity components, suggesting that the decomposition of the connectivity data may be more variable, or more dependent on sample size.

We note some limitations to this work. Firstly, there are a number of considerations when performing structural connectivity analysis: how to parcellate the brain, which algorithm to use for tractography, how to threshold the connectivity matrix, etc., although it remains unclear which methods are the best choice for structural connectivity analyses (Zalesky et al. 2010, 2016) We were limited in our choices in some respect due to the lack of high angular resolution diffusion data. Our method took advantage of state-of-the-art approaches to limit false positives, including the use of anatomical constraints (Smith et al. 2012), we also thresholded our matrices based on a measure of edge weight consistency shown to improve estimate of consistent networks across subjects (Roberts et al. 2017). Secondly, we used a large multisite cohort for this study. Site and scanner variance can impact neuroanatomical measures derived from MRI (Schnack et al. 2010; Fortin et al. 2018). Here, we employed a method for control of batch effects, ComBat, and found that unwanted site variance significantly less apparent within component timecourses (Fig S3). We did not apply this form of site correction to the connectivity data as, although this method had been applied to parametric diffusion maps, e.g.: FA (Fortin et al. 2017), inter-site variation may affect the the initial whole-brain tractography prior to FA weighting. Instead, due to the removal of site variation from the *H* matrix, we only consider edges that vary with each cortical component, and therefore do not vary with site.

In summary, we use a data-driven, multivariate method to identify patterns of regional cortical development that have a hierarchical structure and reliable identified across independent cohorts. We find that these components share developmental trajectories with distinct subnetworks of structural connectivity.

## Acknowledgements

This research was conducted within the Developmental Imaging research group, Murdoch Childrens Research Institute and the Children’s MRI Centre, Royal Children’s Hospital, Melbourne, Victoria. It was supported by the Murdoch Childrens Research Institute, the Royal Children’s Hospital, Department of Paediatrics, The University of Melbourne and the Victorian Government’s Operational Infrastructure Support Program. The project was generously supported by RCH1000, a unique arm of The Royal Children’s Hospital Foundation devoted to raising funds for research at The Royal Children’s Hospital.

Data and/or research tools used in the preparation of this manuscript were obtained and analyzed from the controlled access datasets distributed from the NIMH-supported Research Domain Criteria Database (RDoCdb). RDoCdb is a collaborative informatics system created by the National Institute of Mental Health to store and share data resulting from grants funded through the Research Domain Criteria (RDoC) project. Dataset identifier(s): [DOI: 10.15154/1503353].

