## Supplemental Materials for "Charting shared developmental trajectories of cortical thickness and structural connectivity in childhood and adolescence"

|  | **Table S1: PING demographics by site** | | |  |
| --- | --- | --- | --- | --- |
|  | **Site** | **n** | **age (range)** | **Male** |
|  | a | 87 | 15.37 (3.75-21.00) | 44 (50.6) |
|  | b | 96 | 13.18 (3.58-21.00) | 46 (47.9) |
|  | c | 90 | 13.95 (3.17-20.92) | 45 (50.0) |
|  | d | 93 | 8.99 (3.25-20.25) | 46 (49.5) |
|  | e | 90 | 11.62 (4.17-21.00) | 52 (57.8) |
|  | **Total** | 456 | 12.59 (3.17-21.0) | 233 (51.1) |

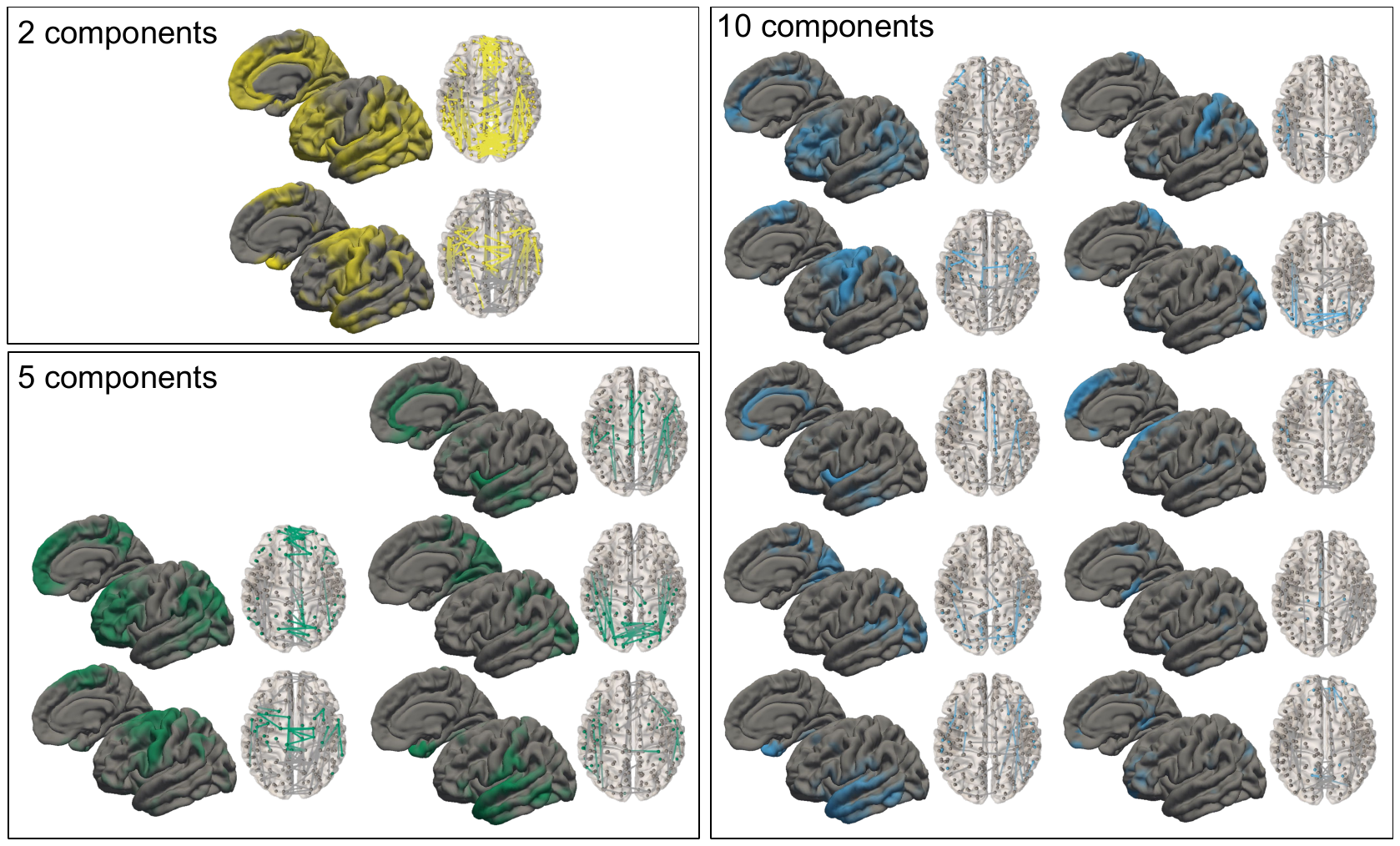
**Figure S1. NMF decompositions into 2, 5 and 10 components.**

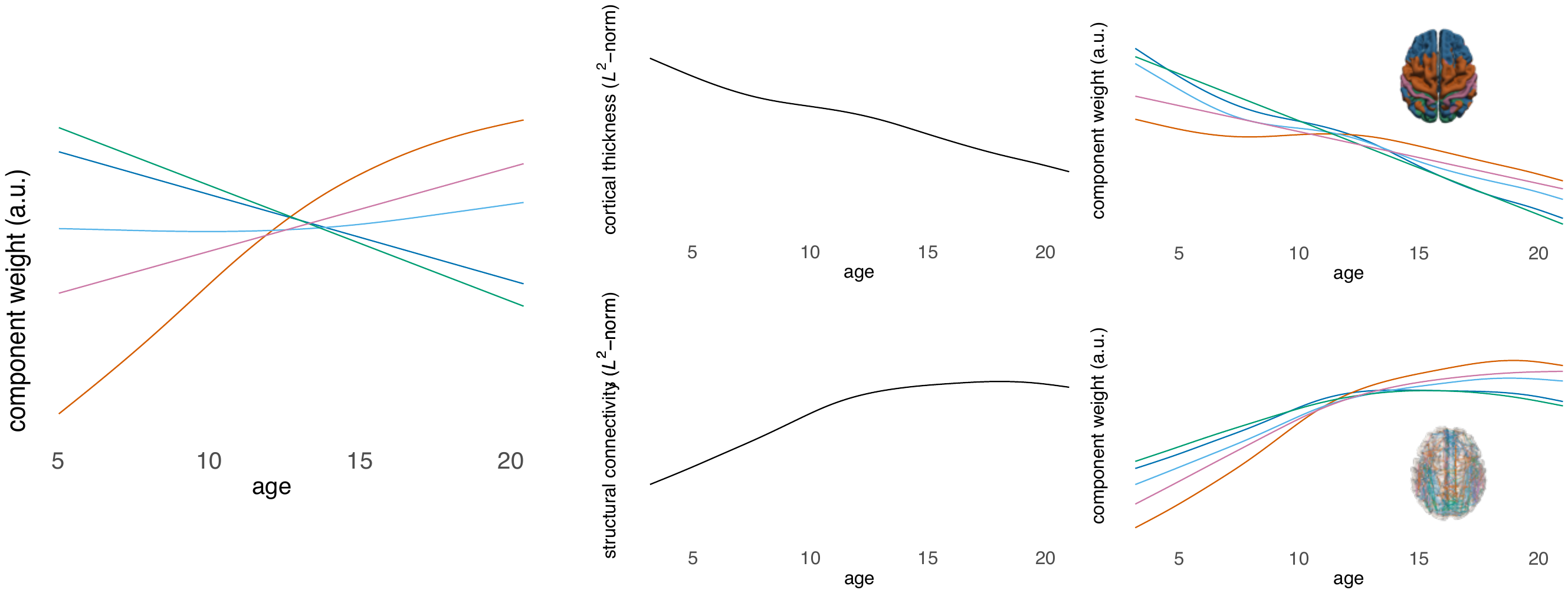

**Figure S2. Normalising component weights.** Components weights for the 5-component solution are shown overlaid on each other (left). Prior to NMF decomposition each subject’s data were normalised to unit norm. The GAM trajectory for the norm of the raw data is shown in the centre column, revealing the mean trend for cortical thickness (top) and connectivity (bottom). Combining the normalised trajectories with the group mean shows the raw trajectories of each component (right)

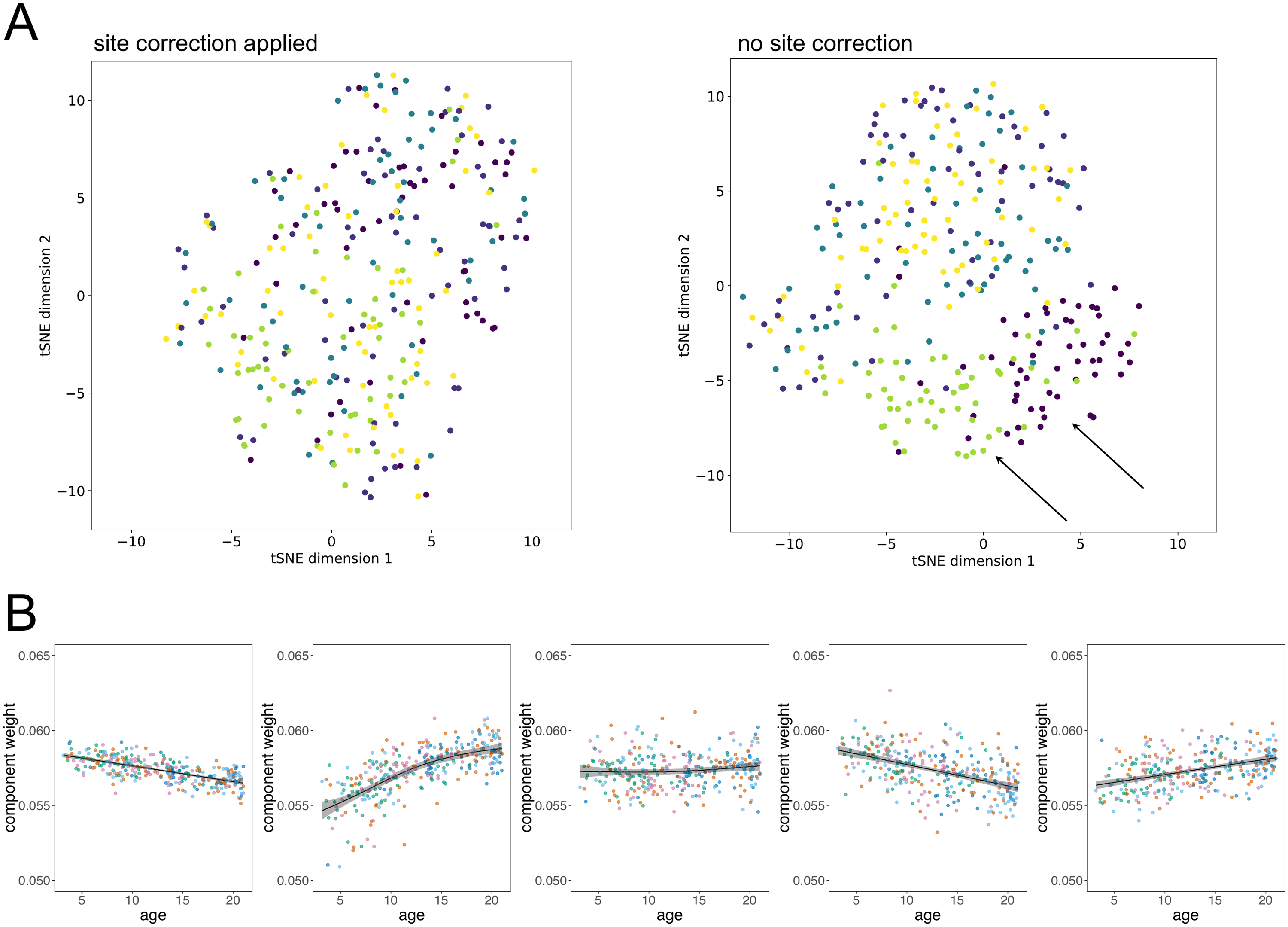

**Figure S3. Removing site variance.** A) Two-dimensional projections of the component timecourses (H matrix) are shown (calculated with UMAP). Without ComBat site correction, significant site bias is present in the component weights (right, arrows), this is not present after site correction (left). B) Component trajectories, as in Figure 2, each datapoint is coloured by site.
